# Biological Filtering and Substrate Promiscuity Prediction for Annotating Untargeted Metabolomics

**DOI:** 10.1101/558973

**Authors:** Neda Hassanpour, Nicholas Alden, Rani Menon, Arul Jayaraman, Kyonbum Lee, Soha Hassoun

## Abstract

Mass spectrometry coupled with chromatography separation techniques provides a powerful platform for untargeted metabolomics. Determining the chemical identities of detected compounds however remains a major challenge. Here, we present a novel computational workflow, termed Expanded Metabolic Model Annotation (EMMA), that aims to strike a balance between discovering previously uncharacterized metabolites and the computational burden of annotation. EMMA engineers a candidate set, a listing of putative chemical identities to be used during annotation, through an expanded metabolic model (EMM). An EMM includes not only canonical substrates and products of enzymes already cataloged in a database through a reference metabolic model, but also metabolites that can form due to substrate promiscuity. EMMA was applied to untargeted LC-MS data collected from cultures of Chinese hamster ovary (CHO) cells and murine cecal microbiota. EMM metabolites matched, on average, to 23.92% of measured masses, providing a > 7-fold increase in the candidate set size when compared to a reference metabolic model. Many metabolites suggested by EMMA are not catalogued in PubChem. For the CHO cell, we experimentally confirmed the presence of 4-hydroxyphenyllactate, a metabolite predicted by EMMA that has not been previously identified as part of CHO cell metabolism.

---

Metabolomics is an expanding field of research that involves the characterization of small molecules in cells, tissues and other biological systems. Metabolites are direct products of enzymatic reactions and provide a functional readout of cellular state ^1–2^. Compared to genes and proteins that are regulated and post-translationally modified, respectively, metabolites are most predictive of the phenotype ^3^. Metabolomics now plays a critical role in many fields including drug discovery and precision medicine, nutritional analysis, and examining environmental responses. Importantly, the ability to collect thousands of measurements on the metabolome using untargeted metabolomics, where thousands of features are measured within the sample under study, and to annotate these measurements with chemical identities, promises to broadly profile the metabolome and revolutionize phenotyping and biological discoveries.

Mass Spectrometry (MS) techniques coupled with liquid or gas chromatography separation techniques, LC-MS or GC-MS, have become standard analytical platforms for untargeted metabolomics ^4^. The LC or GC step aims to separate compounds within the sample, while the MS step ionizes, fragments and detects a fragmentation pattern. There are now techniques for data processing (e.g., peak picking, missing value imputation, and adduct and degenerative feature removal). These tools convert raw MS data into features. Each feature corresponds to an ionized chemical compound, and is characterized using a *spectral signature*, comprising a chromatographic retention time (RT) paired with mass-to-charge ratio (m/z) and relative intensities for the parent compound and its fragments.

Interpretation of metabolomics data is facilitated by assigning putative chemical identities to the features. Relying on the mass of the ionized parent compound for annotation is problematic as a particular mass may be associated with many chemical formulas (e.g., there are 10,132 known molecular structures in PubChem^5^ that are associated with C_20_H_22_N_2_O_4_)^6^. The spectra of detected compounds can be matched against those within an in-house library generated using the same instrument and method as the samples. However, this is impractical due to the large number of compounds detected in an untargeted MS experiment. Instead, feature annotation typically relies on libraries in reference databases. The two largest spectral databases in terms of number of unique compounds, METLIN ^7^ and NIST ^8^, cover less than 17,000 compounds each, which is less than 0.002% of the ~96,000,000 compounds catalogued in PubChem. Due to these limitations, the annotation rate, which we define as the fraction of features annotated with a putative chemical identity, using in-house or spectral databases is typically low. The maximum annotation rate across several metabolomics studies that we surveyed was 16% but averaged only 7.26% ^9–15^.

In recent years, computational tools have become available to recommend a ranked list of chemical structures that best explain a spectral signature. This ranked list is selected amongst a pre-specified *candidate set*, a listing of metabolites with formula weights that match the measured masses of parent compounds in the sample. Earlier tools used rule-based approaches to generate fragmentation patterns of candidate metabolites ^16–18^. Subsequent efforts introduced combinatorial enumeration methods ^19–22^ and machine-learning algorithms. For example, CFM-ID ^23^ uses the candidate set to create a probabilistic model of collision-induced fragmentation process. The model is then used to predict a fragmentation pattern for a given compound. CSI:FingerID ^24^ first predicts a fragmentation tree based on a spectral signature ^25^. CSI:FingerID then uses multiple-kernel learning ^26^ and support vector machines to predict fragmentation tree properties, which are searched against fragmentation tree properties of compounds in a molecular structure database. CSI:FingerID, which outperforms other tools ^19, 23, 27–29^ in terms of accuracy. Despite progress, however, the annotation runtimes, are problematic. For example, annotating 3,868 spectral signatures in GNPS, a public repository of mass spectra data ^30^, using CSI:FingerID against PubChem structures was estimated to take 12 days and 12 hours ^24^. While there are tradeoffs between accuracy and runtime ^24, 31^, the runtime for all tools is dependent on the size of the candidate set. Hence, evaluating candidate sets from large compound databases such as PubChem and ChemSpider remain problematic.

We propose a novel annotation workflow for untargeted metabolomics that addresses current limitations regarding spectral database coverage and computational cost of annotation. The key step is to *filter* the detected masses through a metabolic model that we call an Expanded Metabolic Model (EMM). An EMM includes not only the defined substrates and products of enzymes cataloged for the organism(s) associated with the sample, but also additional metabolites reflecting the potential for promiscuous enzymatic activities. The central premise is that an EMM can be used to define a candidate set that is more comprehensive than a standard genome-scale metabolic model, but still enforces a degree of specificity for the system of interest. Our workflow, termed EMMA (EMM-based Annotation), broadens the search space for annotation beyond compounds in a reference metabolic model assembled from catalog definitions of enzymatic reactions, thus enhancing discovery while avoiding the computational cost of analyzing every compound in large chemical structural databases. We demonstrate the utility of EMMA on untargeted LC-MS data from cultures of Chinese hamster ovary (CHO) cells and bacterial isolates from murine cecum. We compare the candidate sets from reference metabolic models, EMMs, and a large structural database (PubChem). EMMA suggests biologically relevant chemical identities for almost a quarter of measured features, providing a > 7-fold increase in the candidate set size when compared to using a reference metabolic model. Importantly, EMMA allows the discovery of novel relevant putative identities that are not currently catalogued in PubChem. Targeted LC-MS experiments confirmed the presence of a predicted CHO cell metabolite that has previously not been cataloged as a Chinese hamster enzyme substrate or product.

## METHODS

To describe and evaluate the EMMA workflow, we present it alongside two other annotation workflows (**Figure 1**). In describing the workflows, “annotation” refers to the use of any computational annotation tools. A *model-based* annotation workflow (**Figure 1A**)consists of filtering masses of measured metabolites against those expected in the sample based on a metabolic model that is built from a reference genome (or set of reference genomes). Model metabolites with exact masses that match, within a small error, to measured masses are designated as the candidate set. The candidate set is then annotated, where candidates that best explain the experimentally observed spectra are ranked. This workflow provides two advantages. Metabolites within the candidate set are all biologically relevant. Consequently, all computing time will be used to evaluate biologically relevant candidates. While there is now a growing collection of annotated genome sequences (e.g. KEGG database ^32^, BioCyc ^33^, and BiGG ^34^) and tools for the reconstruction of genome-scale metabolic models (GEMs) ^35–36^, the completeness of these models is not guaranteed. GEMs models are typically constructed using sequencing and annotation ^37–38^. Significant experimental and computational efforts are required to augment the models based on gene expression, proteomics, and metabolomics data ^39^. Current models do not account for enzyme promiscuity, where an enzyme transforms alternate substrates in addition to its natural substrate, as defined by a reference metabolic model and as catalogued in organism databases ^40–43^. As a result, defining the candidate set based only on metabolites within the metabolic models naturally limits annotation.

**Figure 1.**
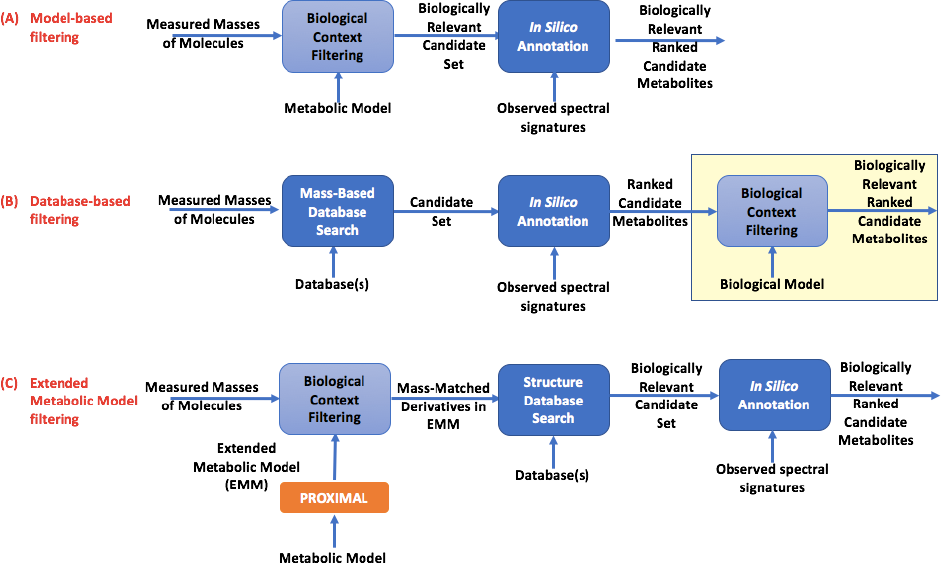
Comparison between annotation workflows. The candidate set for annotation is derived by filtering the measured masses based on: (A) the metabolic model, (B) databases, and (C) extended metabolic model (EMM). The candidate sets in and (C) are biologically relevant, while candidates in (B) prior to filtering may not all be biologically relevant.

Selecting the candidate set from a large database can potentially enhance annotation by increasing the number of measured masses that have a match in the candidate set (**Figure 1B).** This *database-based* workflow first identifies candidate metabolites by querying one or more specified compound databases for all molecules whose exact masses match the experimentally measured masses detected in the sample. The resulting candidate set is then annotated and ranked as in the workflow shown in **Figure 1A**. As the size of the candidate set is large in comparison to the one in the model-based workflow, the annotation runtime increases, and so does the chance of annotation. As annotation accuracy is low, some measurements, however, may be annotated with biologically irrelevant identities, e.g., phytochemicals or drug compounds that cannot possibly accumulate in a mammalian cell culture. The end user could then sift through the ranked candidate metabolites to select biologically relevant candidates. This manual curation is time-consuming and relies on the user’s judgment. Using a metabolic model to filter the ranked candidate metabolites can facilitate this process. However, this results in the same discovery-related limitations as the workflow shown in **Figure 1A,** while also incurring a large computational cost. Importantly, not all the computational cost is necessary. It is highly unlikely that every compound in candidate sets derived from large database is biologically relevant. Using manually curated metabolite databases such as KEGG to derive the candidate set is an attractive option, as the size of the candidate set is reduced when compared to using a large structural database. However, not all biologically relevant compounds are catalogued in such databases.

Our novel annotation workflow (**Figure 1C**), EMMA, applies an EMM-based filter to identify the candidate set. To create this model, we adopt a previously described method, *PROXIMAL* ^44^ (**Supplementary Methods**). From reactant-product pair(s) (RPAIR) of an enzymatic reaction ^45^, *PROXIMAL* identifies a molecular pattern that transforms the reactant into product. Each pattern is associated with a reaction center and its first and second-level neighboring atoms. If a substrate of interest matches a pattern, then the corresponding operator is applied to generate a product, which we call a “derivative” metabolite. The EMM for a system of interest is generated using *PROXIMAL* by applying the operators generated from the enzymatic reactions encoded in the system’s genome(s) to all of metabolites already associated with the system based on the enzymes’ reaction definitions. This generates a set of *derivative* metabolites. The calculated exact masses of derivative metabolites are then used to filter the measured masses. If a derivative has a mass that matches a measured mass, then the KCF or SMILES string of this derivative is searched against a chemical structure database (PubChem) to determine if it has been cataloged with a chemical name and identifier. The masses of metabolites in the reference metabolic model are also matched against the measured masses (as in **Fig 1A**). The union of matched derivatives and reference model metabolites constitute a biologically relevant candidate set. This candidate set is then used for annotation and the candidates are ranked, as in prior workflows. Pseudo-code for the EMMA workflow is provided in **Supplementary Methods**.

## RESULTS

### Datasets, reference metabolic modes, and EMMs

We compared the annotation workflows in **Figure 1** by analyzing untargeted LC-MS data collected on samples from two different biological systems (**Table 1, column group A**). One set of LC-MS experiments were performed on samples from Chinese hamster ovary (CHO) cell cultures grown in a chemically defined medium. The second set of experiments was performed on samples from anaerobic cultures of bacteria collected from murine cecum. Each set of LC-MS experiments comprised two or more different methods. By treating the datasets independently, we were able to explore the influence of sample source and instrument method on EMMA’s performance. Details for the culture and LC-MS experiments are provided in **Supplementary Methods**. The processed data were arranged into feature tables, where each feature was specified by a chromatographic retention time (RT), measured mass (m/z), and a set of associated product ion (fragment) masses and their relative intensities, i.e., MS/MS signature. The reference metabolic models for CHO cell and murine cecal microbiota were derived from genomes in the KEGG database. For the CHO cell, we obtained lists of metabolites and reactions cataloged in KEGG that are associated with the organism code *cge*. The cecal culture is a consortium of many species. We assembled a community-level model based on the taxonomic groups detected in the culture using a previously described procedure ^46^. The numbers of reactions, metabolites and unique masses in the two reference models are listed in **Table 1**(**column groups B**).

**Table 1.** Size of experimental data sets and models. (A) Three experimental datasets under different conditions were collected for the CHO cell, and two for the gut microbiota sample. (B) The size of the metabolic model: number of reactions, metabolites, and unique masses. (C) The size of the expanded metabolic model: number of operators derived using *PROXIMAL*, unique derivatives generated by *PROXIMAL*, unique derivative masses due to *PROXIMAL*. For comparison purposes, the numbers of derivatives and derivative masses exclude those in the metabolic model. (D) Fold increase in number of metabolites and masses when comparing the size of these sets for EMM against the metabolic model.

|  | (A)<br>Experimental data |  |  | (B)<br>Metabolic model |  |  | (C)<br>Expanded metabolic model using<br>PROXIMAL |  |  | (D)<br>Fold change for EMM<br>relative to metabolic model |  |
| --- | --- | --- | --- | --- | --- | --- | --- | --- | --- | --- | --- |
| Biological sample | Dataset | MS mode | Number of measured masses | Number of reactions | Number of metabolites | Number of unique masses | Number of unique operators | Number of unique derivatives | Number of unique derivative masses | Number of metabolites | Number of unique masses |
| CHO cell | HilNeg | negative | 2,502 | 1,619 | 1,353 | 775 | 2,392 | 76,745 | 17,930 | 56.72 | 23.14 |
|  | HilPos | positive | 3,856 |  |  |  |  |  |  |  |  |
|  | SynNeg | negative | 5,336 |  |  |  |  |  |  |  |  |
| gut microbiota | Neg | negative | 1,651 | 1,381 | 1,307 | 779 | 2,756 | 94,186 | 23,356 | 72.06 | 29.98 |
|  | Pos | positive | 1,657 |  |  |  |  |  |  |  |  |

The EMM for each sample was generated using biotransformation operators derived from each model (**Table 1, column group C**). EMMs augment a metabolic model to include molecules that are not originally part of the metabolic model. This augmentation increases the number of unique masses within the model. The number of biologically relevant molecules in the candidate set thus significantly increases (**Table 1, column group D**) when compared to the number of metabolites in the reference metabolic model (57x and 72x for CHO cell and the gut microbiota, respectively). Similarly, the number of unique masses in EMM is increased over the number of unique masses in the reference metabolic model (23x and 30x for the CHO cell and the gut microbiota, respectively). EMMs thus promise to provide a large annotation space when compared to the reference metabolic model.

### Novel Annotation opportunities

Compared to using the reference metabolic model for a biological sample, using an EMM as the search space during metabolite annotation increases the size of the candidate set for annotation in terms of: (a) matching to a larger number of measured masses, and (b) suggesting a larger set of putative chemical identities. Using these two metrics, we compare the size of the “biologically relevant candidate sets” in the model- and EMMA-based workflows and compare that with the size of the candidate set using PubChem (**Table 2**). A small percentage of the measured masses are matched to masses of metabolites in the metabolic model. On average, 3.31% of measured masses can be potentially annotated using the metabolic model only. When using the EMMs, this number increases to 23.92%, a 7.6-fold increase. When restricting the EMM derivatives to those that have a catalog entry in PubChem, the annotation rate drops to 5.12% as there are many compounds that are not yet catalogued in PubChem, currently the largest structural database. Using PubChem, the number of mass matches are in the millions. Not all such metabolites are biologically relevant. The use of reference metabolic models or large database such as PubChem therefore provide some limitations in annotation when compared to using EMMs. Using EMMA allows for novel biological discovery by suggesting biologically relevant compounds not in PubChem and reduces the annotation space considerably.

**Table 2.**
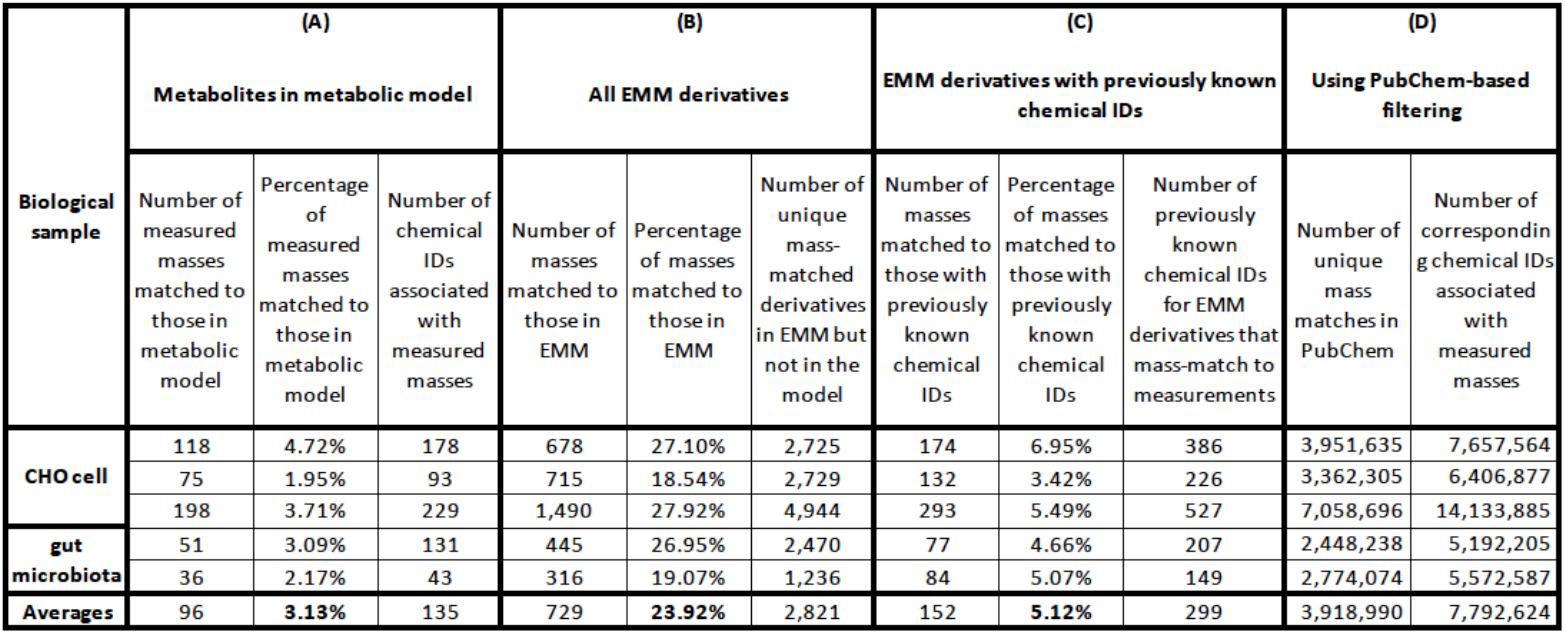
Candidate set size using different workflows. (A) Candidate set size when using the model: number of measured masses that match to metabolites in the model, the equivalent percentage of the number of measured masses reported for experimental data in **Table 1**, and corresponding number of chemical identities. (B) Candidate set size when using EMMA: number of measured masses that match to metabolites in the EMM, equivalent percentage in reference to the number of measured masses reported for experimental data in **Table 1**, and corresponding number of chemical identities. (C) Further filtering of the EMM derivatives reported in column group (B) to include only mass measurements that match to previously known chemical IDs as reported in PubChem, and reporting the number of matched masses, the relative percentage of these masses to the number of measured masses reported for experimental data in **Table 1**, and the corresponding number of chemical IDs. (D) Size of the candidate set when filtering using PubChem.

As the size of the KEGG database is significantly smaller than PubChem, KEGG may not be as computationally prohibitive to use for annotation. Further, using only a biological database such as KEGG for annotation guarantees biological relevance of candidate metabolites. A question that often arises is regarding the benefits of utilizing a general database for annotation compared to when employing a database that mostly comprises biomolecules. Using the EMMA workflow as a reference and restricting derivatives to those with chemical identities in PubChem, we are able to explore and quantify the benefits. Specifically, we utilized the EMMA workflow to identify candidates sets for our datasets. We then compared the EMM candidates against those obtained using the database-based workflows using KEGG and PubChem (**Table 3**). Many candidate molecules identified by EMMA that are catalogued in KEGG (e.g. For CHO cell HilNeg data, 93 out of 174 candidate compounds). However, there were also EMMA candidate compounds found in PubChem that are not catalogued in KEGG. For our datasets and using EMM metabolites as a reference, there is at least 2x or more additional biologically relevant candidates in PubChem for each candidate identified in KEGG. The 2x-fold increase over KEGG is a lower bound on the number of biologically relevant metabolites in PubChem. Using a large database such as PubChem thus significantly increases biologically relevant annotation opportunities when compared to KEGG. Relying only on small biological databases limits annotation. EMMA provides an alternative candidate set that provides different tradeoffs between annotation opportunities and speed.

**Table 3.**
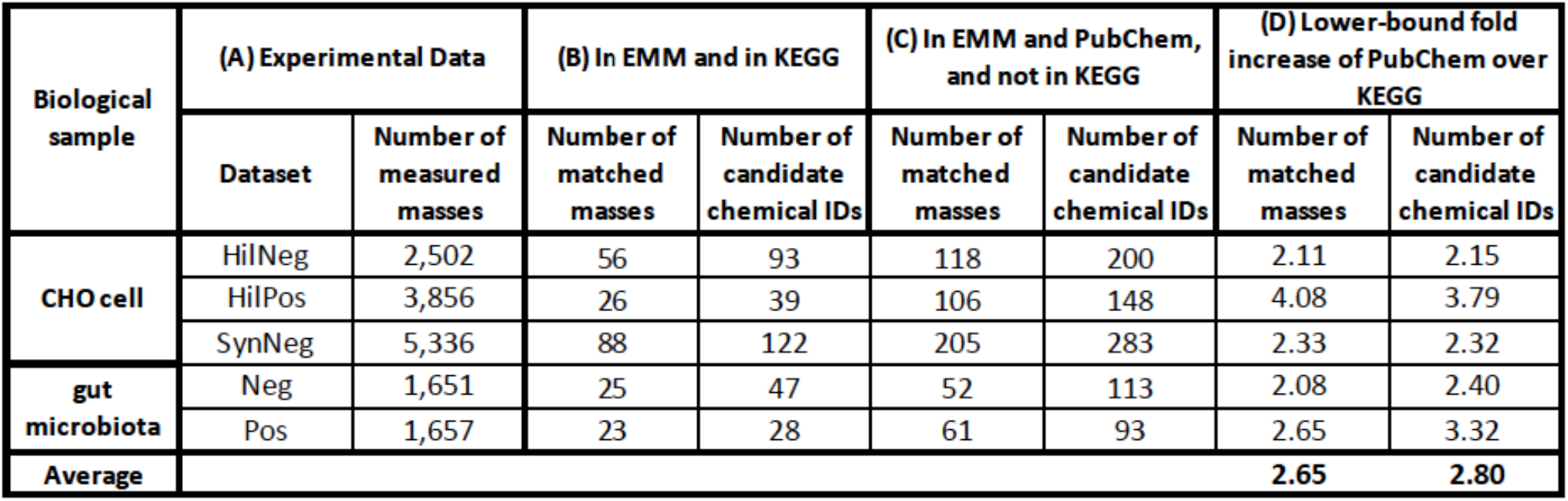
Using EMMs to compare the annotation opportunities of PubChem against KEGG. (A) Experimental data for different data sets (repeated for convenience). (B) Number of matched masses and candidate chemicals found using EMMA that are reported in KEGG. (C) Number of matched masses and candidate chemicals found using EMMA reported in PubChem but not in KEGG. (D) Lower-bounds on discovery of biologically relevant matched masses and candidate chemicals when using PubChem over KEGG.

### Computational time required for annotation

To generate the candidate set as the input to *in silico* annotation analysis in database-based workflow (**Figure 1B**), we identified metabolites in the KEGG and PubChem databases that mass-matched within 10 ppm to the masses in our experimental data for each dataset. We investigate the computational time required to annotate the candidate sets from EMMs and from the combined PubChem and KEGG databases. Annotation of each candidate set identified by EMMA required a handful of hours, averaging 2.5 hours per data set (**Table S1, group A**). The number of candidate metabolites from databases PubChem and KEGG for each of our datasets exceeded 5 million candidates, with an average dataset size of 7.8 million candidates (**Table S1, group B**). It was computationally prohibitive to annotate all mass-matched metabolites from the databases. To calculate the required runtime, we estimate it using annotation runtimes based on the EMMA workflow (**Table S1, group A**). Dividing the runtime by number of metabolites in the candidate set, on average, annotation requires 0.0085 hours per match. Using this average, the estimated runtime for annotation of database-based workflow is computed for each dataset. The average required runtime per dataset is over 65,000 hours (**Table S1, group B**).

### Experimental validation of EMMA

We next investigated whether any of the derivatives predicted by EMMA and matched to a detected MS feature based on mass and MS/MS signature could be experimentally confirmed with a chemical standard. We selected eight predicted derivatives that matched an LC-MS feature for CHO cell samples (**Table 4, group A**). The selection was based on two factors: the rank assigned by the *in silico* annotation tool and availability from a vendor. The selected derivatives are: salicylaldehyde, one of the three isomers of hydroxybenzaldehyde; 4-hydroxyphenyllactate, a tyrosine metabolite; acetoacetamide, a monocarboxylic acid amide of acetoacetic acid; 5-aminopentanoate, a lysine degradation product; glutarate, produced in lysine and tryptophan metabolism; 3-methoxyanthranilate, an ester of anthranilic acid; 2-hydroxyphenylacetic acid, associated with styrene degradation pathway; and 4-pyridoxate, a product of vitamin B_6_. When using KEGG as a database for annotation, CFM-ID ranked 6 of the 8 derivatives as the highest ranked candidates, while two of the derivatives were not in KEGG (**Table 4, group B**). Further, a small number of candidate matches were found for each mass measurement. When using PubChem as a database for annotation, all derivatives ranked among the three top candidates (**Table 4**, **group C**). As expected, the number of putative matches increased when compared to the number of matches using KEGG. We analyzed the number of reactions in CHO that contributed an operator that was used to generate each derivative and the number of E.C. numbers that are associated with each set of reactions (**Table 4, group D**). The number of reactions and enzymes varied for each derivative. For example, 12 different reactions catalyzed by 15 enzymes correspond to the operator that generated 4-hydroxyphenyllactate, whereas only one reaction and enzyme corresponds to the operator that generated acetoacetamide.

**Table 4.** Right candidate metabolites identified by EMMA used for experimental validation. (A) Candidate mass and name. (B) Ranking of metabolite and number of candidates that matched mass measurement using KEGG. (C) Ranking of metabolite and number of candidates that matched mass measurement using PubChem. (D) The number of reactions that yielded the *PROXIMAL* operator that yielded each candidate metabolite and the associated number of enzymes that catalyze these reactions. (E) The status of experimental validation

| (A)<br>Candidate metabolites |  | (B)<br>KEGG |  | (C)<br>PubChem |  | (D)<br>PROXIMAL |  | (E) |
| --- | --- | --- | --- | --- | --- | --- | --- | --- |
| Mass measurement (Daltons) | Candidate metabolite identified by EMMA | Rank | Matches | Rank | Matches | Number of Reactions used to derive operator | Number of E.C.s. associated with reactions | Experimentally validated? |
| 122.04 | Salicylaldehyde | 1 | 1 | 1 | 1 | 1 | 1 | No |
| 182.06 | 4-Hydroxyphenyllactate | 1 | 2 | 1 | 4 | 12 | 15 | Yes |
| 101.05 | Acetoacetamide | 1 | 1 | 2 | 3 | 1 | 1 | No |
| 117.79 | 5-Aminopentanoate | 1 | 2 | 1 | 5 | 4 | 4 | No |
| 132.04 | Glutarate | 1 | 1 | 3 | 6 | 12 | 11 | No |
| 167.06 | 3-Methoxyanthranilate | 1 | 1 | 2 | 3 | 8 | 2 | No |
| 152.05 | 2-Hydroxyphenylacetic acid | NA | 1 | 1 | 4 | 1 | 1 | No |
| 183.05 | 4-Pyridoxate | NA | 0 | 1 | 1 | 1 | 1 | No |

Comparing the RTs and MS/MS spectra of standards for these chemicals against the corresponding CHO cell culture sample features (**Supplementary Methods**), we were able to confirm correct annotation of 4-hydroxyphenyllactate (**Figure 2**). This demonstrates that the EMMA can indeed identify a novel metabolite that is not found among metabolites cataloged for the organism of interest, in this case the Chinese hamster. In addition to KEGG, we searched for 4-hydroxyphenyllactate in BioCyc and MetaCyc. Neither database associated this metabolite with the Chinese hamser. In KEGG, 4-hydroxyphenyllactate is associated with three enzymatic reactions. Reactions 4-hydroxyphenyllactate:NAD+ oxidoreductase (R03336) and 4-Hydroxyphenyllactate:NADP+ oxidoreductase (R03338) are both catalyzed by D-hydrogenase (E.C. 1.1.1.222, which was recently deleted and transferred to E.C. 1.1.1.110) and hydroxyphenylpyruvate reductase (E.C. 1.1.1.237). Reaction 3-(4-hydroxyphenyl)lactate hydro-lyase (4-coumarate-forming) (R08766) is associated with an enzyme that have yet to be characterized (E.C. 4.2.1.-). It is unlikely that the source of 4-hydroxyphenyllactate in our sample is exogenous, as our cell culture medium was chemically defined and did not include this metabolite. Further evidence that the metabolite is endogenously derived is provided by a recently updated GEM for the CHO cell in the BiGG database ^47^. This GEM reconstruction (iCHOv1) included 4-hydroxyphenyllactate as a “universal” metabolite that could be formed enzymatically, but for which a specific gene encoding the enzyme remains unknown.

**Figure 2.**
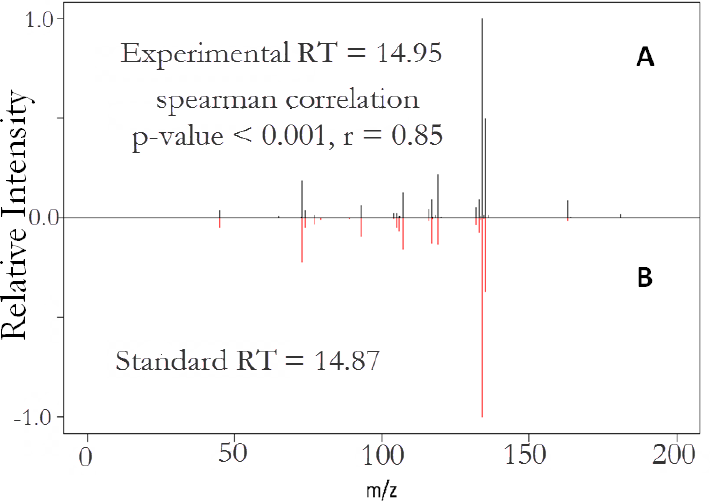
Mirror plot for 4-hydroxyphenyllactate, KEGG compound C03672. (A) experimental data collected using untargeted metabolomics from the CHO cell culture. (B) data from high-purity chemical standard. This is considered a match by retention time (RT difference < 3 minutes) and by MS/MS (spearman rank correlation p-value < 0.05 and r-value > 0.6)

## DISCUSSION AND CONCLUSION

Our EMMA workflow addresses the challenge of creating an annotation candidate set that is enzymatically relevant to the sample under study and that includes metabolites beyond what is already catalogued in reference metabolic models. One important contribution of the work is conceptually separating the engineering of the candidate set from annotation. Only a limited number of prior works engineered candidate sets and in limited ways. These works focused on exclusions of particular elements, substructures or compounds ^31^ or on inclusion sets ^48^. Filtering PubChem compounds using PubMed Medical Subject Heading (MeSH) labels ^49^ can reduce the candidate set size by including only naturally occurring compounds that are biologically relevant (e.g. carbohydrates, lipids, etc.). However, only a tiny fraction (124,049 compounds) of PubChem compounds has MeSH labels. Filtering using current MeSH labels was reported to reduce a candidate set of 62,782 structures that match in mass to 3,868 compounds in GNPS dataset to only 36 compounds ^50^.

Results from comparing the three workflows emphasize the need for optimizing the engineering of candidate sets. We demonstrated for our two case studies that using candidate sets from large databases is computationally prohibitive, as others have also noted ^24^. We also demonstrated that using a biological database such as KEGG yields a small candidate set when compared to using a large structural database. We further demonstrated that using a reference metabolic model is inadequate as only a very small percentage (3.31% on average) of measurements can be annotated. In this regard, EMMA contributes two key advances. First, filtering candidate chemicals using an EMM allows for the identification of novel metabolites that are missing from a GEM reconstruction. This advance addresses the need to enable discovery, which is inherently limited in the simpler approach of using a model comprising only known metabolites to filter the candidate chemicals or when using a small biological database, without incurring a prohibitive computational cost. Second, filtering the measurements through an EMM specific for the system of interest provides a biologically relevant and computationally feasible candidate set. This advance eliminates unnecessary and time-consuming computations on chemicals from large databases that are likely irrelevant to the system of interest. Not all biologically relevant candidates from a large database are in the EMM. This issue could be addressed by further expansion of the EMM candidate set by the repeated application of the biotransformation operators derived from the reference model to derived promiscuous products. While we demonstrated the EMMA workflow using specific tools and databases (e.g., CFM-ID for annotation; PubChem and KEGG databases) the overall workflow is generic and can be readily modified to use other *in silico* annotation tools.

EMMA relies on a reference metabolic model for annotation. Other recent studies have also exploited the metabolic network to enhance annotation. One method, iMet, suggests that neighboring metabolites within a metabolic network have similar MS/MS spectra and trains a classifier to predict if two spectra belong to neighboring metabolites ^51^. The classifier is trained using MS/MS spectra from spectral databases and mass differences between reactant pairs from KEGG that are not specific to the biological sample. Another method, BioCAN creates a network based on measured features and assigns aggregate annotation scores based on spectral lookups and annotation tools^52^. Mummichog maps measured features to metabolic models, and performs statistical pathway and module enrichment ^53^. There are also other studies that exploit putative biotransformation for annotation. In one method, the mass difference between a pair of features is matched against mass differences between substrate-product pairs of common metabolic conversions (e.g., oxygenation, acetylation, etc.), with a match indicating a potential biochemical transformation between the pair of detected feature masses ^54^. These transformations can be used to propagate metabolite annotation from an identified metabolite to its potential reactants and products. In contrast to this method, EMMA does not require any MS/MS training data and utilizes biological context that is specific to the sample. There is a common limitation when using metabolic models to improve annotation. Genome-scale metabolic reconstructions can be incomplete, especially for non-model organisms. Our biotransformation prediction by PROXIMAL suggests that the metabolite 4-hydroxyphenyllactate may result from the promiscuous activity of one or more carboxylic acid dehydrogenases expressed in the CHO cell on the substrate molecule 4-hydroxyphenylpyruvate.

This work presents the first *in vivo* experimental evidence for a computationally predicted metabolite derived through promiscuous action of an enzyme. Using a chemical standard, we confirm the presence of 4-hydroxyphenyllactate in a CHO cell culture, even though there is no documented gene associated with the CHO cell metabolism that can catalyze the reaction with 4-hydroxyphenyllactate as product. We were however able to confirm only one out of the eight predicted metabolites. This could be due to inaccuracies in the rankings by the annotation tool. For example, CSI:FingerID reports an accuracy of only 39.5% when annotating a data set from MassBank by searching PubChem. The low confirmation rate can also be due to the assumption that all enzymes are promiscuous. As an enhancement, we are currently investigating methods to improve *PROXIMAL* to rank predicted derivatives based on enzyme designations as generalists or specialists ^55^ and participation in primary or secondary metabolism ^56^.

Despite limitations due to the underlying potentially incomplete metabolic models and to the accuracy of current annotation tools, EMMA demonstrates utility in creating an expanded, biologically relevant candidate set and in utilizing it to enhance annotation. Importantly, EMMA promises to offer annotation opportunities beyond those possible with metabolic models without the high computational cost of searching large structural databases that contain many non-biological compounds.

## AUTHOR INFORMATION

### Notes

The authors declare no competing financial interest.

### Author Contributions

S.H. conceived the EMMA workflow. N.H. performed all the computational work. K.L. conceived the experimental validation. K.L.’s lab provided the experimental data for CHO. N.A. conducted the experimental validation for the CHO Cell. A.J.’s lab provided the experimental data. R.M. conducted the experimental work on the murine cecal cultures. N.H., K.L. and S.H. edited the manuscript with input from N.A.

## ACKNOWLEDGMENT

This work was supported by the National Institute of Health under project number 1R03CA211839-01 and NSF grant #1421972.

## ASSOCIATED CONTENT

### Supporting Information

Supplementary Methods is a PDF file that provides a detailed description of PROXIMAL and EMMA.

