## Supplementary Methods for "Biological Filtering and Substrate Promiscuity Prediction for Annotating Untargeted Metabolomics"

### 1. Computational Methods

#### 1.1 Identifying biologically relevant molecules beyond those in the metabolic model

The sample's metabolic model can be augmented into an expanded metabolic model based on enzyme promiscuity. To this end, we generalized the pattern matching method described in our earlier work, *PROXIMAL*, which was originally developed for identifying possible bio-transformation products of xenobiotic chemicals in the liver due to Cytochrome P450 (CYP) enzymes. The key idea in *PROXIMAL* is to approximate enzyme activities through bio-transformation operators that act on molecular fragments. To expand the metabolic model, each bio-transformation operator is applied to each metabolite within the model.

The bio-transformation operators are constructed as follows. The transformation of each fragment is be specified using Reaction Center, Difference Region, and Matched Region (RDM) patterns [1]. The RDM patterns of metabolic enzymes are available from the KEGG reaction pair (RPAIR) database [1], and specify local regions of similarities/differences for reactant-product pairs based on chemical structure [2]. An RDM pattern consists of three parts: a Reaction Center (R) atom that exists in both the substrate and reactant molecule on the boundary between Matched and Non-Matched Regions, Difference Region (D) atoms that are adjacent to the R atom but also part of the Non-Matched Region, and Matched Region (M) atoms adjacent to the R atom in the Matched Region. A lookup table is constructed based on the RDM patterns of enzymes associated with reactions in the model. The "key" in the lookup table consists of the R and M atom(s) and adjacent neighbors in the reactant, while the "value" represents the R and D atom(s) in the product. For each potential R pattern matched in the query molecule, a set of transformations are looked up in the table and applied to the query molecule.

To illustrate how *PROXIMAL* functions, an example is shown in **Fig.** S**1**. In **Fig. S1A**, a specific reversible reaction (KEGG reaction ID: R03534) transforms 2-oxoglutarate (KEGG compound ID: C00026) to 2-hydroxyglutarate (KEGG compound ID: C02630). The reactant and product molecules are encoded using KEGG atom types [2], while the atom numbers, extracted from KEGG KCF files, are specified in parenthesis following the type of atoms in the structure of each compound. Each reactant-product atom pair is then entered into a transformation table (**Fig. S1B**). The transformation table identifies patterns of change in atom types along with a local context through the transformation of reactant to product. To identify transformation patterns, *PROXIMAL* aligns the atoms in reactant-product structures, and adds each atom in reactant and its corresponding atom in the product as a new row to the transformation table. The ordering of the rows in the table is determined by the ordering of atoms in the reactant molecule structure (**Fig. S1B**). Having the transformation table, any reactant atom that is aligned to a product atom with a different type will be considered as a potential reaction center. In this example, rows 1 and 4 demonstrate two potential reaction centers in reactant compound: C5a and O5a. To add specificity to these transformations, the lookup table keys are augmented to include two-level nearest neighbors including the reaction center (**Fig. S1C**). To visualize the concept of two-level nearest neighbors, we used a color code in **Fig. S1A** illustrating this concept for one of the potential reaction centers, O5a. The potential reaction center O5a is shown in red. The first-level neighbor (adjacent neighbor) C5a is shown in blue, and the second-level neighbors (distant neighbors) C1b and C6a are shown in green. The same biotransformation can be derived by multiple reactions cataloged in KEGG. For this specific example, reactions with KEGG IDs R00267, R00342, R00709, R01000, R01388, R01392, R01394, R01513, R03104, R03688, and R07136 can lead to the same bio-transformation pattern. Similarly, the set of adjacent and distant neighbor atoms for the potential reaction center C5a can be extracted (**Fig. S1C**). The set of distant neighbors always include the reaction center.

Given a query compound, *PROXIMAL* applies a select set of transformations from the lookup tables at one or more matching sites, or reaction centers, of the query compound, where several derivatives are possible (**Fig. S2**). Considering each atom in the query molecule as a potential reaction center, *PROXIMAL* creates a neighbors table containing list of adjacent and distant neighbors for each of the potential reaction centers. *PROXIMAL* then looks for matches between the generated list and keys in the lookup table. In a case of match, *PROXIMAL* applies the matched key's value to the reaction center and its neighbors to generate a product. Query compound 4-hydroxyphenylpyruvate (KEGG compound ID: C01179) is demonstrated with atom types in **Fig. S2A**. For each atom in the structure of the query compound, a list of adjacent and distant neighbors is generated and added to neighbors table (**Fig. S2B**). Comparing the neighbors table against the keys in the lookup table (**Fig. S1C**) shows row 4 of the neighbor table, with potential reaction center O5a, as a match. Application of the matched key's value to the reaction center and its neighbors leads to a biotransformation product 4-hydroxyphenyllactate with KEGG compound ID: C03672 (**Fig. S2C**).

To create an EMM given a reference catalogued metabolic model, one or more operators are derived from substrate-product pairs associated with each reaction. Operators are then applied to all metabolites within the model. The expanded model size depends on the number of operators and metabolites of the reference model.

**A**

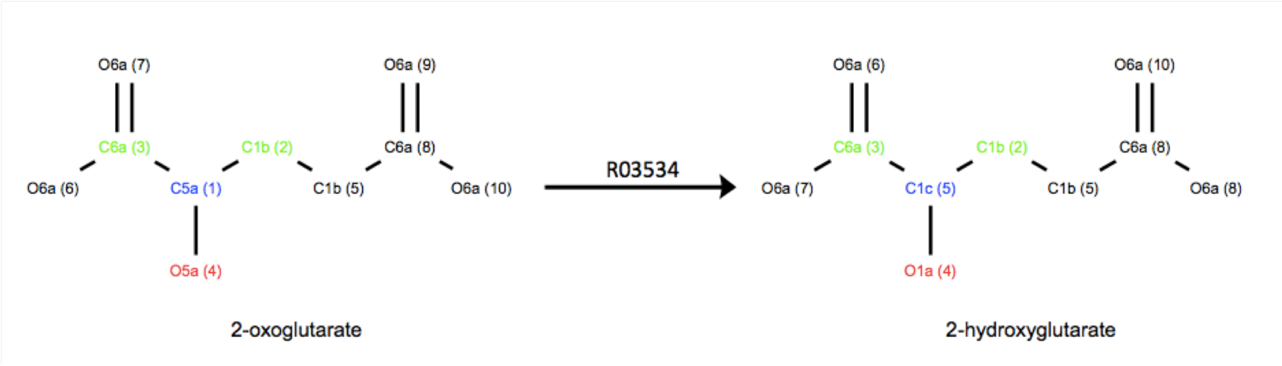


**B**

| Reactant | | Product | |
|---|---|---|---|
| Atom # | Atom type | Atom # | Atom type |
| 1 | C5a | 1 | C1c |
| 2 | C1b | 2 | C1b |
| 3 | C6a | 3 | C6a |
| 4 | O5a | 4 | O1a |
| 5 | C1b | 5 | C1b |
| 6 | O6a | 7 | O6a |
| 7 | O6a | 6 | O6a |
| 8 | C6a | 8 | C6a |
| 9 | O6a | 10 | O6a |
| 10 | O6a | 9 | O6a |

**C**

Keys

| Reaction center | Adjacent neighbor 1 | Distant neighbors 1 | Adjacent neighbor 2 | Distant neighbors 2 | Adjacent neighbor 3 | Distant neighbors 3 |
|---|---|---|---|---|---|---|
| C5a | O5a | C5a | C6a | C5a, O6a, O6a | C1b | C5a, C1b |
| O5a | C5a | O5a, C6a, C1b | --- * | --- | --- | --- |

* Not applicable

Values

| Reaction center | Adjacent neighbor | Added functional group |
|---|---|---|
| C1c | O1a, C6a, C1b | --- * |
| O1a | C1c | --- |

* Not applicable

**Fig. S1.** Illustration of generating lookup tables by *PROXIMAL*. (A) Reactant and product of an enzymatic reaction R03534, for which *PROXIMAL* aims to derive possible corresponding bio-transformations (operators). (B) Transformation table containing matching atom pairs in reactant and product compounds. (C) Potential operators: key table specifies the transformed substructure in reactant. Value table specifies the modification in product corresponding to the content of key table.

**A**

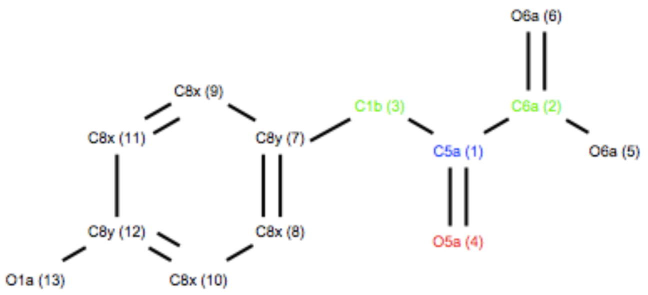


4-hydroxyphenylpyruvate

**B**

| Atom # | Reaction center | Adjacent neighbor 1 | Distant neighbors 1 | Adjacent neighbor 2 | Distant neighbors 2 | Adjacent neighbor 3 | Distant neighbors 3 |
|---|---|---|---|---|---|---|---|
| 1 | C5a | C6a | C5a, O6a, O6a | C1b | C8y, C5a | O5a | C5a |
| 2 | C6a | C5a | C6a, C1b, O5a | O6a | C6a | O6a | C6a |
| 3 | C1b | C8y | C1b, C8x, C8x | C5a | C1b, C6a, O5a | --- * | --- |
| 4 | O5a | C5a | O5a, C6a, C1b | --- | --- | --- | --- |
| 5 | O6a | C6a | O6a, O6a, C5a | --- | --- | --- | --- |
| 6 | O6a | C6a | O6a, O6a, C5a | --- | --- | --- | --- |
| 7 | C8y | C1b | C8y, C5a | C8x | C8y, C8x | C8x | C8y, C8x |
| 8 | C8x | C8y | C8x, C1b, C8x | C8x | C8x, C8y | --- | --- |
| 9 | C8x | C8y | C8x, C8x, C1b | C8x | C8x, C8y | --- | --- |
| 10 | C8x | C8x | C8x, C8y | C8y | C8x, C8x, O1a | --- | |
| 11 | C8x | C8x | C8x, C8y | C8y | C8x, C8x, O1a | --- | --- |
| 12 | C8y | C8x | C8x, C8y | C8x | C8y, C8x | O1a | C8y |
| 13 | O1a | C8y | O1a, C8x, C8x | --- | --- | --- | --- |

*Not applicable

**C**

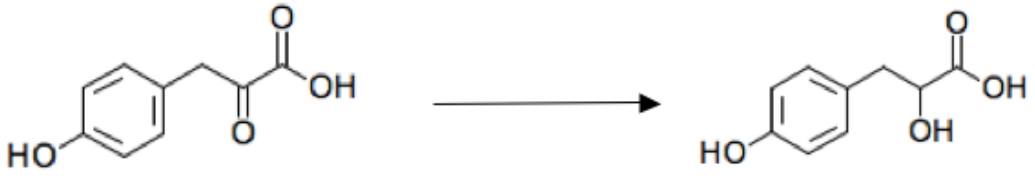


4-hydroxyphenylpyruvate 4-hydroxyphenyllactate

**Fig. S2.** Illustration of application of lookup table to a query molecule by *PROXIMAL* to generate the potential bio-transformation products. (A) A query compound represented by KEGG atom types. (B) Table of neighbors generated considering each atom in the query compound as a potential reaction center. Row 4 in the generated table matches to one of the keys in the lookup table shown in Fig. S1C. (C) The product 4-hydroxyphenyllactate is the result of applying the matched key's value to the query compound.

### 1.2 Details of the EMMA Annotation workflow

---

EMMA workflow

---

**Procedure** EMMA (**in** *metabolic model*, **in** *measured masses of molecules*, **in** *observed Spectral signatures*, **in** *database(s)*, **out** *biologically relevant ranked candidate metabolites*)

```
Begin
    1. use model reactions in metabolic model to generate biotransformation lookup tables
    2. identify mass-matched derivatives in extended metabolic model (EMM)
    for each metabolite in metabolic model
        2a. apply biotransformation lookup tables on metabolite to generate potential
        derivatives
        for each derivative in potential derivatives
            2b. calculate, M, the mass of derivative
            for each mass measurement m in measured masses of molecules
                2c. use an error margin to generate a mass interval
                if M falls into mass interval
                    add derivative to mass-matched derivatives in EMM
                end if
            end for
        end for
    end for

    3. compare mass-matched derivatives in EMM to database(s), add the ones that match
    structurally to a metabolite in a database into biologically relevant candidate set

    4. use an in silico fragmentation tool to score biologically relevant candidate set against
    observed spectral signatures and output biologically relevant ranked candidate metabolites
end
```

**Fig. S3.** Pseudo code of the EMMA workflow

Given a model (list of metabolites and reactions) as well as tandem MS data (mass measurements of parent molecules and associated spectral signatures) for a biological sample, the goal is to associate each mass measurement with a compound ID. The workflow of EMMA, **Fig. 1** (main document), is outlined in **Fig. S3**.

In step 1, *PROXIMAL* is used to create transformation lookup tables based on enzymatic reactions in the input model. In step 2a, the biotransformation information stored in the created lookup tables is applied to model metabolites to generate a set of potential derivatives in EMM. In step 2b, the monoisotopic masses of atoms are used to calculate the mass of each potential derivative. In step 2c, the calculated masses are compared within the specified error margin against measured masses to generate a list of mass-matched derivatives in EMM. In step 3, the mass-matched derivatives in EMM are structurally compared against compound databases to add structurally-matched metabolites to the list of biologically relevant candidate set. In step 4, biologically relevant candidate set metabolites are scored and ranked against the observed spectral signatures using *in silico* fragmentation leading to generate biologically relevant ranked candidate metabolites. We chose to use 10 PPM mass error margin as MetFrag [3] in implementation of EMMA workflow. We used CFM-ID [4] as the fragmentation prediction tool for scoring the candidate metabolites.

#### 1.3 Curating metabolic models for CHO

The model for the CHO cell was curated from the KEGG database as follows. A list of pathways associated with the CHO cell is selected, excluding pathways with numbers larger pathway number than 1100 as the most main metabolic pathways have numbers less than 1100. For each reaction in the pathways, the reactant and product compound IDs, RPAIRS, and enzyme IDs were retrieved.

#### 1.3 Computational time required for annotation using each workflow

The table below provides detailed information regarding the runtime of the EMMA vs database-based workflow.

**Table S1**. Computational speed up of EMMA workflow over database-based workflow using PubChem and KEGG for our datasets. (A) For the EMMA workflow, the following data is provided for each dataset: candidate set size generated by EMMA, relevant CFM-ID runtime for EMMA to perform annotation on the candidate set, and average runtime per match. (B) For the database-based workflow, the size of the candidate and the estimated run time of CFM-ID is provided.

| Biological sample | Dataset | (A) EMMA workflow | | | (B) Database-based workflow | |
|---|---|---|---|---|---|---|
| | | size of candidate set | runtime (hrs) | average CFM-ID runtime per match, EMMA(h) | size of candidate set | estimated runtime (hrs) |
| CHO cell | HilNeg | 386 | 3.5748 | 0.00926114 | 7,657,564 | 70,917.77 |
| | HilPos | 226 | 1.5375 | 0.006803097 | 6,406,877 | 43,586.61 |
| | SynNeg | 527 | 4.0425 | 0.007670778 | 14,133,885 | 108,417.89 |
| gut microbiota | Neg | 207 | 2.021 | 0.009763285 | 5,192,205 | 50,692.98 |
| | Pos | 149 | 1.343 | 0.009013423 | 5,572,587 | 50,228.08 |
| Averages | | 299 | 2.50 | 0.0085 | 7,792,624 | 64,768.67 |

### 2 Materials and methods for collecting data from untargeted metabolomics and experimental validation

#### 2.1 CHO Cell Culture

Chinese Hamster Ovary (CHO) cells expressing recombinant monoclonal antibody were cultivated using proprietary, chemically defined media and feed (Biogen Idec, Cambridge, MA) as described in a previous study [5] in bioreactors that controlled process parameters including temperature, dissolved oxygen concentration and pH. Samples removed from the bioreactor were centrifuged to pellet cells and the supernatant was gently aspirated and stored in a fresh vial at -80°C.

#### 2.2 CHO Sample Extraction

CHO supernatant samples were thawed on ice and added to extraction solvent (100% methanol) at a sample to methanol ratio (v/v) of 1 to 3 and vortexed for 15 seconds. Protein was precipitated and removed by centrifugation at 4°C and 15,000 x g for 15 minutes. 200μL supernatant was carefully aspirated and transferred to a fresh sample vial. The samples were concentrated by first drying using a vacuum concentrator (Eppendorf Vacufuge 5301) and reconstitution in 100μL methanol/water (50/50 v/v).

**2.3 Cecal Cultures**

Whole cecum was taken from eight weeks old female C57BL/6J mice (Jackson Laboratories, ME) maintained on an ad libitum chow diet (8604 Teklad Rodent diet). Mice were euthanized using asphyxiation with CO2 and excised cecum was transported to an anaerobic chamber (Coy chamber) in an anaerobic transport medium (Anaerobic systems). Inside the chamber, cecal contents were extracted and made into a slurry using pre-reduced PBS containing 0.1% cysteine. 1% of this slurry was inoculated into 10 ml of Gut microbiota medium [6]. Samples were stored at -80°C for metabolites extraction.

**2.4 Cecal Sample Extraction**

Metabolites were extracted for liquid chromatography-mass spectrometry (LC-MS) from 5 ml of the culture samples. Samples were mixed with 5 ml of cold methanol and 2.5 ml of chloroform and homogenized using lysing matrix E beads (MO BIO, CA) on a bead beater (VWR, PA). The samples were homogenized for one min, cooled on ice for one minute, and homogenized again for another 2 min. The samples were then centrifuged at 10,000g at 4 °C for 10 min. The supernatant was filtered through a 70-μm sterile nylon cell strainer into a clean sample tube and mixed with 0.6 ml of ice-cold water using a vortexer. This mixture was centrifuged again at 10,000g for 5 min to obtain phase separation. The upper and lower phases were separately collected using a syringe while taking care not to disturb the interface. The upper phase was dried to a pellet using a vacufuge (Eppendorf, Hauppauge, NY), and stored at −80 °C until further analysis. Prior to LC-MS analysis, the dried samples were reconstituted in 50 μl of methanol/water (1:1, v/v).

**2.3 Untargeted LC-MS**

Untargeted analysis was carried out on a quadrupole time-of-flight (qTOF) mass spectrometer (TripleTOF 5600+, AB Sciex) with an electrospray ionization source in tandem with a binary pump HPLC system (1260 Infinity, Agilent). To maximize the number of metabolites that could be detected, the mass spectrometer was run in both positive and negative ionization modes and two separate liquid chromatography columns were used (hydrophilic interaction and reverse phase). The mass spectrometer was operated in information dependent acquisition mode (IDA) to collect MS/MS fragmentation data for as many detected masses as possible. The IDA experiment consisted of a TOF MS survey scan and four dependent product ion scans (MS/MS) for molecules that met set criteria. Dynamic background subtraction was used to limit redundant MS/MS collection and maximize quality by only selecting masses that had intensities that rose quickly over several scans.

Two different reverse phase columns were used depending on the sample. For the CHO samples, a polar endcapped C18 column (Synergi Hydro-RP) was used. For the microbiota samples, a polar embedded C18 column (Synergi Fusion-RP) was used. Both columns used identical mobile phases and gradients. For these columns, Solvent A was 0.1% formic acid in water and Solvent B was

0.1% formic acid in methanol. The column was maintained at 15°C with a 55 minute gradient elution with the following set points: t = 0 – Solvent B = 3%, t=8 – Solvent B = 3%, t = 38 – Solvent B = 95%, t = 45 – Solvent B = 95%, t = 47 – Solvent B = 3% and t = 55 – Solvent B = 3%.

For hydrophilic retention, an aminopropyl column was used (Luna $NH_2$, Phenomenex). Solvent A was 20 mM ammonium acetate in 95:5 water:acetonitrile pH adjusted to 9.45 using ammonium hydroxide and Solvent B was 100% acetonitrile. The column was maintained at 25°C with a 60 minute gradient elution with the following set points: t = 0 – Solvent B = 85%, t=15 – Solvent B = 0%, t = 28 – Solvent B = 0%, t = 30 – Solvent B = 85% and t = 60 – Solvent B = 85%.

#### 2.4 Data Preprocessing

Data preprocessing was performed using custom R scripts implementing xcms [7] [8] [9] and CAMERA [10].Peak-picking and alignment was performed using xcms to generate a feature table of m/z, RT pairs. CAMERA was used to identify and remove isotopes, in-source fragments and adducts.

#### 2.5 Experimental Validation

For the eight compounds that were selected for experimental validation, high-purity standards were ordered from Sigma-Aldritch (St. Louis, MO). Experimental verification was carried out using the same LC-MS method used to analyze the CHO cell samples. An assignment was considered a match if it was confirmed by at least two orthogonal methods including monoisotopic mass, retention time and MS/MS fragmentation pattern [11]. A match was made using retention time if the retention times of the experimental sample and standard were within 1 minute of each other. An MS/MS match was confirmed using a Spearman Rank Correlation performed on shared fragment peaks between the sample and standard with a p-value less than 0.05 and R value greater than 0.6.
